# Pitch-size crossmodal correspondence in tortoises (*Testudo hermanni*)

**DOI:** 10.1101/2023.01.31.526484

**Authors:** Maria Loconsole, Gionata Stancher, Elisabetta Versace

## Abstract

Humans spontaneously match information coming from different senses, what we call crossmodal associations. For instance, high pitch sounds are preferentially associated with small objects, and low pitch sounds with larger ones. While previous studies reported crossmodal associations in mammalian species, evidence for other taxa is scarce, hindering an evolutionary understanding of this phenomenon. Here we provide evidence of pitch-size correspondence in a reptile, the tortoise *Testudo hermanni*. Tortoises showed a spontaneous preference to associate a small disc (i.e., visual information about size) to a high pitch sound (i.e., auditory information), and a larger disc to a low pitch sound. These results suggest that crossmodal associations may be an evolutionary ancient phenomenon, potentially an organising principle of the vertebrate brain.

## 1. Introduction

The ability to combine information obtained through different senses is crucial for perceiving the environment and responding to different situations. Humans display systematic crossmodal associations between different sensory inputs that could belong to a same stimulus [1]. For instance, both adults [2,3] and infants (6-months-old [4], 30-months-old [5], but not 4-months-old [4]) preferentially match high pitch sounds with visually smaller objects, and low pitch sounds with larger ones. This would suggest that different sensory modalities are connected, and sensory inputs that are more likely to come from the same distal stimulus are processed together. In the natural environment, there are systematic correspondences between features encoded in different sensory modalities. For instance, sources of illumination often come from above, hence it is more likely that objects in elevated spatial positions would be brighter than those in lower spatial positions [6,7]; similarly, smaller animals are more likely to produce high pitch sounds, while larger animals are more likely to produce low pitch sounds, due to the anatomical structure of their vocal tract [8,9]. For this reason, the ability to make crossmodal associations could confer and advantage to the individual in making accurate predictions, reducing uncertainty and allowing the formation of coherent and meaningful representations of objects and events [10,11]. A comparative perspective could provide important insights on the phylogenetic origin of crossmodal associations, by investigating whether species from different taxa could benefit from the same facilitations.

The pitch-size crossmodal association has been reported in mammalian species but is yet to be investigated outside of the mammalian clade. Wild chimpanzees seem to modulate pitch-size associations, producing high-pitch or low-pitch vocalizations to signal smaller or larger trees, respectively [12]. Domestic dogs spontaneously pair abiotic high- and low- pitch sounds with artificial objects of different sizes [13]. This association may have an adaptive value because the pitch of vocalizations can be used to identify the body size of conspecifics [14]. Human co-domestication might have also played a role, as the same effect was found in human participants when asked to match a picture of a dog (having a small or large body size) with its growl (high or low in pitch) [9]. To investigate the presence of crossmodal pitch-size association in a far related taxon, here we explore whether a reptile, the tortoise (*Testudo hermanni*), can associate pitch and size, using abiotic artificial stimuli such as playbacks of pure high vs low pitch sounds and yellow disks of different sizes.

Tortoises can process auditory information (e.g., they can locate an auditory source to escape a maze [15] and respond to unexpected auditory stimuli, for instance by retracting the head in the shell [16]). To the best of our knowledge, there is no complete analysis for sound perception in tortoises, but studies on *Testudo hermanni* have shown that this species responds to airborne sounds in the range of 10 to 940 Hz [16,17]. Copulating sounds emitted by males can influence females’ choice of partners, and indicate male fitness [17,18]. Although male Hermann tortoises process pitch information, there is no evidence suggesting that they have a spontaneous preference for high or low pitch sounds. Similar to what had been described for humans [2,4,5] and dogs [13], and in line with the idea that crossmodal associations correspond to real-world feature associations, in the present study we hypothesized to find a spontaneous preference to match a high pitch sound with a small shape, and a low pitch sound with a larger shape.

## 2. Materials and methods

### 2.1. Subjects and rearing conditions

We tested 10 adult male tortoises (*Testudo hermanni*). Tortoises were housed in group in an outdoor environment at *Sperimentarea* (Rovereto Civic Museum Foundation, Italy). Subjects that entered the test were kept in isolation within comfortable fences and individually monitored until completion of test. They had free access to fresh water and received food during the experimental procedure (strawberries) and in the housing environment (lettuce and herbs). Experiments were run in July, the period of highest activity for these animals (tortoises fall into hibernation approximately from November to April).

### 2.2. Apparatus

The arena consisted of an 80×30×40 cm high corridor made of wood. This was divided in three separate areas, a central area (30×30 cm) used as the animal’s starting point, and two choice areas (25×30 cm) accessible from the central area by a flap door that could be pushed open by the tortoise (**Fig. 1A**). On each of the two short sides of the arena, we located a loudspeaker. The arena was centrally lit from above (~150 cm) through a 400 W halogen lamp.

**Fig. 1.**
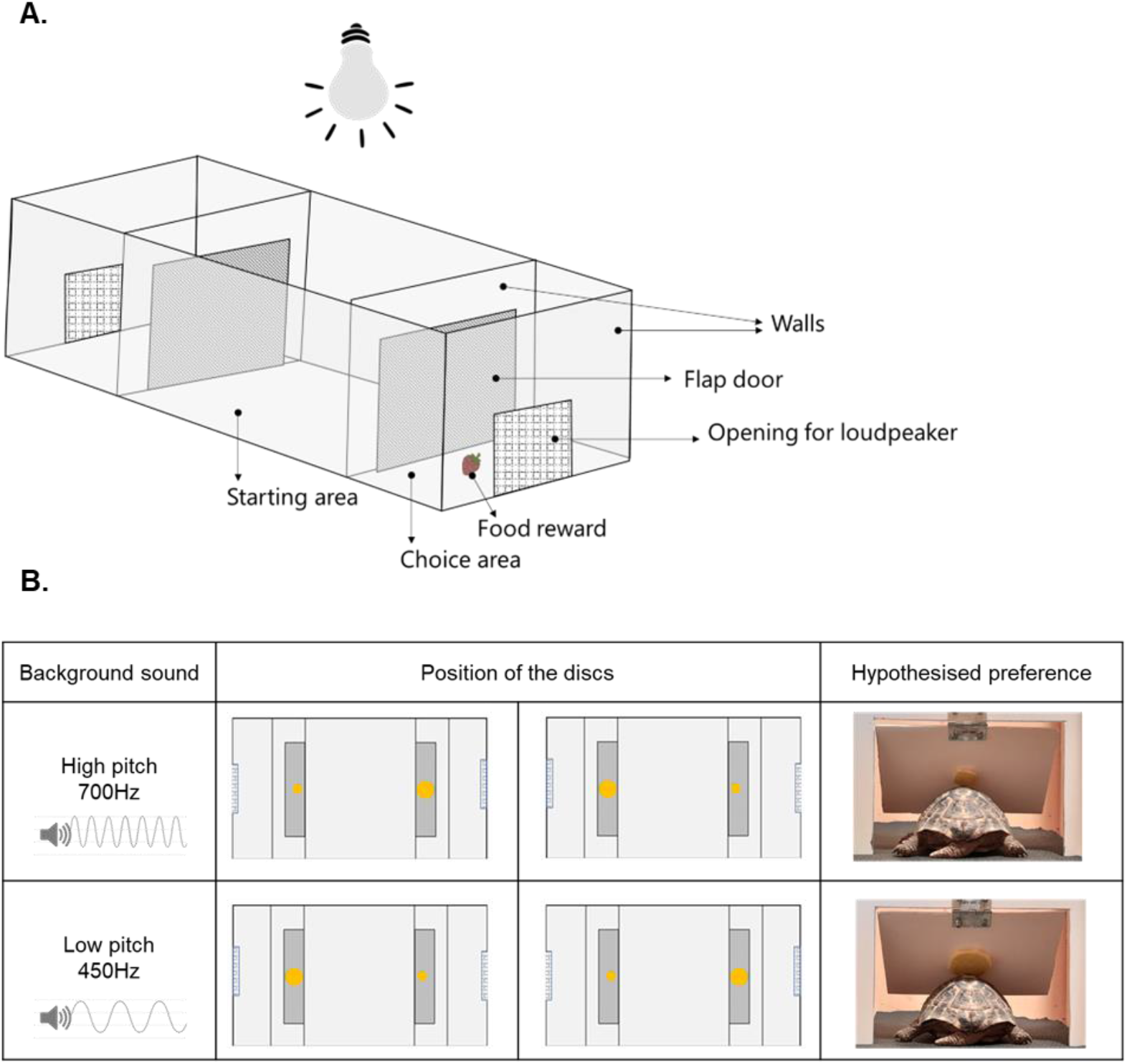
A. The experimental arena: At the beginning of each trial, the tortoise was placed in the centre of the arena (starting area). The tortoise could move from the centre to an adjacent side area (choice area) via the flap door placed on either side. In each trial, a food reward (i.e., a piece of strawberry) was located on the side area signalled by the playing loudspeaker. A small opening on the edge of the arena hosted the loudspeaker. **B. Test**: During test, tortoises received no reward. Both speakers played the same pitch sound (either both high, or both low pitch). Each flapping door could be identified by a yellow disc placed on its centre, either a small, or a large one. The position of the discs was counterbalanced between trials. We hypothesised a preference for the door with the smaller disc in the high-pitch sound condition and a preference for the door with the larger disc in the low-pitch sound condition.

### 2.3. Training

Tortoises were trained to follow an auditory cue to locate the correct area where to enter. The audio track alternated low and high pitch sounds (450 and 700 Hz, ~65 dB) each lasting ten seconds (**S1**). A small piece of strawberry (0.5×0.5 cm) was placed in front of the playing speaker as reward. Tortoises were trained to push the flap door with their foreleg to enter one of the choice areas via a shaping procedure in which the doors were gradually closed until the desired behaviour was learnt. Each subject moved to the following step of the shaping procedure when responding correctly to six consecutive trials, within three minutes per each trial.

### 2.4. Test

Tortoises were tested in a free choice task in two testing blocks of 12 consecutive trials each, with no food reward. During test, both speakers played either the high pitch or the low pitch sound (alternated between trials). One flap door presented a larger yellow disc (5 cm diameter, 0.5 cm thickness), and the other door presented a smaller yellow disc (2.5 cm diameter, 0.5 cm thickness) in the centre. The right/left position of the larger disc, and the audio (high, or low pitch) were randomized between trials (**Fig. 1B**). We scored which door the tortoise chose (i.e., the one with the larger or the smaller disc), considering a choice complete when the subject passed the door with its head and at least half of the shell.

### 2.5. Data analysis

We analysed data using the statistical software R 4.0.2 [19], using a generalized linear mixed-effect model (R package: lme4 [20]) with a binomial structure, the dependent variable being dichotomous (i.e., smaller or larger disc), and including as independent variables the pitch (high or low) and the testing block (first 12 trials, second 12 trials). We included the testing block as a relevant factor to monitor whether performance would change due to loss of motivation (i.e., being the test unrewarded, animals might stop responding to the testing stimuli and behave at chance after some trials) or learning (i.e., animals might modify their behaviour following a prolonged exposure to the testing stimuli) [21].

We run a post-hoc analysis with Bonferroni correction to determine the direction of the effect of the predictors (R package: emmeans [22]). Graphs were generated using ggplot2 [23]. Alpha was set to 0.05.

## 3. Results

In line with our hypothesis, we found an interaction between pitch and block (*X*^2^ = 8.199, p = 0.004) (**Fig. 2**), with a significant crossmodal association present in the first block but not in the second. In the first block (i.e., first 12 trials), when comparing the probability of choosing the smaller disc in the high pitch sound (P(small) = 0.672, SE = 0.058) or the low pitch sound (P(small) = 0.377, SE = 0.067) conditions, we found that tortoises performance significantly differed between the two conditions (contrast (high/low): odds ratio = 3.383, SE = 1.307, z = 3.154, p = 0.009), with animals preferring the small disc when hearing the high pitch sound, and the large disc when hearing the low pitch sound. In the second block (i.e., last 12 trials), when comparing the probability of choosing the smaller disc in the high pitch sound (P(small) = 0.481, SE = 0.07) or the low pitch sound (P(small) = 0.56, SE = 0.061) conditions, tortoises didn’t exhibit a significant preference (contrast (high/low): odds ratio = 0.731, SE = 0.27, z = −0.847, p = 0.832).

**Fig. 2.**
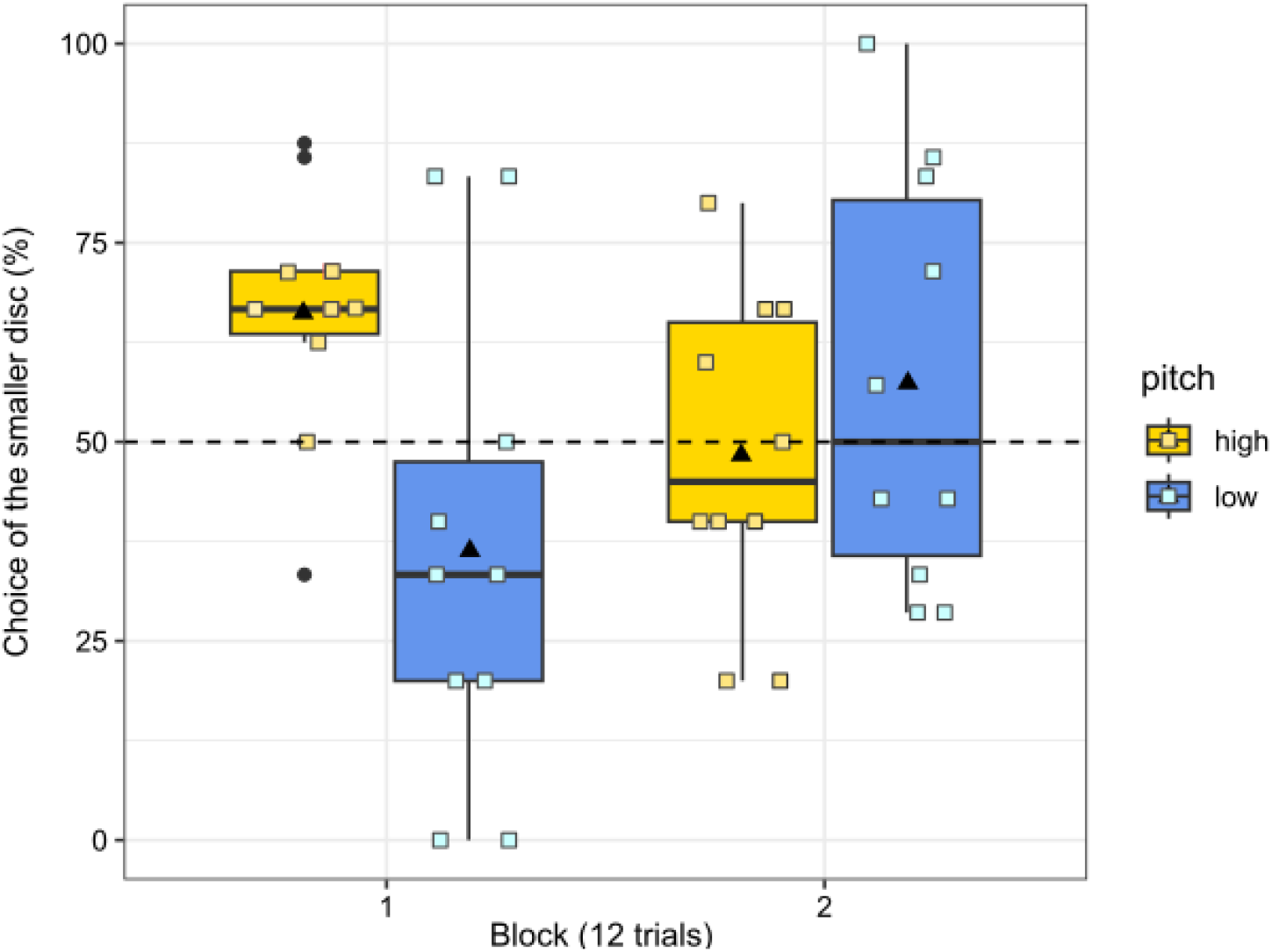
Results. Percentage of choice for the smaller disc for trials presenting high (yellow) vs low (blue) pitch. The dashed line indicates chance level (50%). In each boxplot, the black triangle indicates the mean, and the horizontal bar indicates the median. The black dots indicate the outliers. The squares indicate the performance of each individual subject.

## 4. Discussion

This study investigated pitch-size crossmodal correspondences in adult tortoises (*Testudo hermanni*), focusing on males. We described a spontaneous association between the two sensory dimensions of pitch (high vs low) and size (small vs large), similar to what previously reported in humans [2–5] and domestic dogs [13]. These results indicate that crossmodal associations between acoustic pitch and visual size of objects might be a generalized and predisposed perceptual phenomenon, common to different taxa. A predisposition for matching certain dimensions of the stimuli could be beneficial for the individual when interacting with the environment, facilitating certain associations while hindering those that are less frequent. In the case of the pitch-size correspondence there are congruent statistical regularities in nature. For instance, the size of an animal influences the pitch of its calls, due to the anatomical structure of the vocal chords (a larger vocal apparatus resonates at lower frequencies [8]). As such, there is an advantage in spontaneously associating a feature in one modality (e.g., acoustic pitch), predicted on the basis of a feature encountered in a different modality (e.g., visual size). Previous studies suggested that crossmodal associations may be a shared characteristic in mammals, possibly reflecting a similar pre-natal organization and development of the involved neural mechanisms [10,12,13]. In light of the new evidence reported in our work, and recent data from an avian model [21,24], we suggest that there is a far more ancient origin of crossmodal associations, as these can be observed in phylogenetically distant species such as birds and reptiles. In particular, instances of spontaneous crossmodal associations have been recently identified at the onset of life, in chicks tested soon after hatching, suggesting that the brain may be spontaneously organised to allow crossmodal predictions/associations [24]. It is yet to be determined to what extent relevant experiences play a role in determining the emergence of certain associations [25].

Our study shows that the spontaneous association between pitch and size fades after repeated trials. This is unlikely the result of fatigue, as test never exceeded 20 minutes, and the cognitive load was low (i.e., spontaneous choice between two alternatives). A possible interpretation could be that crossmodal associations serve as an initial strategy to quickly respond to novel situation where the animal is subject to multiple stimulations from different sensory modalities [21]. Hence, to serve their scope, crossmodal associations should not be rigid, but rather be subject to habituation or learning, adapting to the different situations that the animals is facing similar to other predispositions [26,27].

Overall, our data provide evidence of pitch-size crossmodal associations in a reptile. This constitutes an essential step in developing our understanding of the evolutionary origins of crossmodal associations, revealing commonalities in the perceptual organisation of phylogenetically far related species. Based on our discovery, further research can determine how crossmodal associations are formed, whether they fully depend on predispositions, and to what extent can be modified in response to individual experiences and exposure to environmental statistical regularities.

## Supporting information

Supplemental S1 experimental stimuli

## Ethics

This study complied with all applicable national and European laws concerning the use of animals in research. All the employed were examined and approved by the Rovereto Civic Museum Foundation Ethical Review Committee (approval no. 0000304 dd 11/07/22).

## Data accessibility

The dataset generated during the experiment is available as supplementary material **S2**.

## Authors’ contribution

M.L.: Conceptualization, Methodology, Validation, Formal analysis, Investigation, Writing – Original Draft, Visualization; G.S.: Methodology, Resources, Writing – Review & Editing, Supervision, Project administration; E.V.: Conceptualization, Methodology, Resources, Writing – Review & Editing, Supervision, Project administration.

## Acknowledgements

We wish to thank all the staff from *Sperimentarea* for the help in building the experimental arenas and animal care, and the Rovereto Civic Museum Foundation for providing the facilities to carry out this research. We thank Federico Fracchetti and Marco Papiccio for their help with data collection and animal care.

